# Upper limb rehabilitation after stroke: constraint versus intensive training. A longitudinal case-control study correlating motor performance with fMRI data

**DOI:** 10.1101/2022.12.22.521566

**Authors:** Soline Bellaiche, Danielle Ibarolla, Jérôme Redouté, Jean-Christophe Comte, Béatrice Medée, Lisette Arsenault, Audrey Mayel, Patrice Revol, Ludovic Delporte, François Cotton, Gilles Rode, Yves Rossetti, Dominique Boisson, Maude Beaudoin-Gobert, Jacques Luauté

**Author notes:** **Correspondence:** Maude Beaudoin-Gobert.

## Abstract

**Background:** The reproducible beneficial effect of constraint-induced movement therapy (CIMT) in hemiparetic stroke patients makes it a good model to study brain plasticity during rehabilitation procedures.

**Objective:** Assess the functional brain reorganization induced by each of the two components of CIMT: (i) non-affected upper-limb constraint and (ii) intensive training of the paretic arm.

**Methods:** Brain activity of a right hemiparetic chronic stroke patient and of 10 healthy controls was recorded with a functional magnetic resonance imaging (fMRI) during a finger opposition task. For the patient, a total of 8 assessments were performed, before and after each component of CIMT. At each time point, brain activity during movement was compared with rest. Patient’s results were first compared to the control group and then correlated to motor performance across sessions.

**Results:** Constraint-therapy-related improvement was correlated with a decrease of cerebral activity in sensory-motor regions of both the affected and the non-affected hemispheres. Intensive-therapy-related improvement was correlated with the recruitment of pre-motor cortices and cerebellum in both hemispheres.

**Conclusions:** Two different patterns of brain activity underlie the effects of intensive training and constraint which could account for the respective effect of each component of the therapy.

## 1 Introduction

The Constraint-Induced Movement Therapy (CIMT) is a rehabilitation technique based on two theoretical principles (1): (i) overcome the learned non-use of the paretic limb by constraining the movement of the less affected limb, (ii) Facilitate motor recovery through intensive training of the paretic limb. The concept of acquired ‘non-use’ was initially studied in primates after experiments with deafferentation of the upper limb (1). The deafferented monkey no longer used its affected limb even though it was not paralyzed. Restraint of the healthy limb was followed by reuse of the affected limb. The concept of acquired non-use arose from these observations. It has been hypothesized that, in the acute phase of the deficit, the animal experiences the uselessness of the affected limb or even its dangerousness. This observation leads to a functional exclusion of this limb. Despite the recovery that may occur after the acute phase, this limb is still considered by the animal as “useless”. If the healthy limb is constrained, the animal experiences the possibilities of using the affected limb and uses it again in daily life. To investigate the cortical reorganization underlying these phenomenon, Nudo and colleagues (2) mapped cortical territories in deaffereanted monkeys using intracortical microstimulation before the lesion and after the rehabilitation protocol. More specifically, they demonstrated an extension of the representation of the fingers to the detriment of the proximal segments after the period of intensive rehabilitation. These results led to the proposal of restraining the healthy upper limb in the hemiplegic patient to stimulate the reuse of the paretic limb. Indeed, Taub et al (3) developed a rehabilitation protocol to try to improve the prehension abilities of the hemiplegic patient after a stroke. This protocol consists of 2 types of management during a two-week period: (i) restraint of the healthy upper limb for 90% of the time awake for 14 days; (ii) intensive rehabilitation of the paretic upper limb 6 hours per day, 5 days per week, combining physical therapy and daily activities (eating, throwing a ball, playing dominoes…).

Thus, CIMT has been shown to improve significantly upper-limb function in a few proportions of hemiparetic patients after stroke (around 6% of stroke patients) that have recovered at least 10° of wrist and finger extension (4). Several reasons make CIMT a good model to study the functional brain reorganization induced by a technique of upper-limb rehabilitation in hemiparetic patients after stroke: (i) the program shares the same two component framework: constraint of the less affected limb and intensive rehabilitation of the paretic limb, which are both theoretically well-based (ii) CIMT could show reproducible beneficial effects on the upper-limb function (5); and, (iii) compared to control interventions of equal duration and dose, CIMT yields greater improvements in a variety of indicators of upper-limb function in adult survivors of a stroke with residual upper-limb movement (6).

Functional imaging (Positron Emission Tomography, PET or functional Magnetic Resonance Imaging, fMRI) allows studying the reorganization of brain activity −so-called vicariation-during a specific motor task (for a review see (4)). Most studies performed until now have compared brain activity before vs. after CIMT (for a review see (8)). Except for correlation analyses between behavioral changes and brain activity, these comparisons should be interpreted with caution because, in a classical factorial design, an observed difference in brain activity does not necessarily prove a causal link with the effect of rehabilitation. Brain activity changes may be also interpreted as a test-retest effect.

Two models of post-stroke reorganization have been proposed, namely the vicariation *versus* the interhemispheric competition models. For instance, the role of the ipsilateral (contralesional) motor cortex remains debated and generates a significant interest within the field of neurosciences. The vicariation model predicts that the contralesional motor cortex reflects an adaptative phenomenon whereby contralesional motor regions are recruited to generate a new motor output to the affected hand (for a review see (9)). Alternatively, according to the interhemispheric competition model, the activation of the contralesional motor cortex has been shown to favor an imbalance of interhemispheric inhibition, deleterious for hand recovery (10). Identifying these phenomena to manipulate them by a rehabilitation method, like CIMT, in order to provide clinical benefits for the patient is a crucial issue in terms of neuro-rehabilitation. However, previous functional imaging studies did not show consistent results (11–21)(Table 1).

**Table 1:** CIMT and ipsilateral motor cortex (ILMC) activation: review of the current literature. “Ipsilateral” refers to the hand used to perform the task. Rehabilitation procedure: Classical refers to Wolf et al. 2006 and comprises a constraint of the non-affected arm 90% waking hours; intensive therapy of the affected arm 6 hours per day during 5 sessions per week for a 2 weeks period. C: constraint - IT: intensive training - F: flexion - E: extension - CIMT: constraint-induced movement therapy

| References | Number of patients | Delay since stroke onset | Task (fMRI) | Rehabilitation procedure | ILMC |
| --- | --- | --- | --- | --- | --- |
| Levy et al. 2001 | 2 | > 4 mos | Finger tapping | Classical | Increased activation in one patient after CIMT |
| Johansen-Berg et al. 2002a | 7 | > 6 mos | Finger F/E | C: 90% waking hours<br>IT: 1 h/d<br>2 wks | No correlation with motor improvement |
| Schaechter et al. 2002 | 4 | > 6 mos | Finger F/E | C: waking hours<br>IT: 4 hrs/d<br>5 sessions/wk; 2 wks | Increased activation after CIMT in two patients out of three interpretable |
| Dong et al. 2006 | 8 | > 3 mos | Pinch (finger 1 vs. fingers 2 et 3) | Classical | Decreased activation after CIMT but important variability between subjects |
| Hamzei et al. 2006 | 6 | > 1 yr | Wrist F/E | Classical | No change |
| Szaflarsk et al. 2006 | 4 | > 1 yr | Finger tapping | C: 5 hrs/d; IT: 30 mns<br>3 sessions/wk; 10 wks | Increased activation in 3 patients |
| Sheng & Lin 2009 | 1 | 4 mos | Finger tapping | Classical | Increased activation after CIMT |
| Lin et al. 2010 | 5 | > 3 mos | Finger F/E | C: 6 hrs/d; IT: 2 hrs/d<br>5 sessions/wk; 3 wks | Increased activation after CIMT |
| Zhao et al. 2012 | 12 | > 4 mos | Finger F/E | C: 8 hrs/d; IT: 5 hrs/d<br>5 sessions/wk; 3 wks | Increased activation after CIMT compared to before CIMT in 6 out of 12 patients |
| Könönen et al. 2012 | 11 | > 7 mos | Finger F/E | C: ?; IT: 6 hrs/d<br>5 sessions/wk; 2 wks | activation increase in voxels associated to CIMT in 8 out of 11 patients |
| Stark et al. 2012 | 2 | > 4 mos | Wrist, elbow, ankle movements | Classical | New activation appeared in both patients after CIMT |
| Laible et al. 2012 | 7 | > 1 yr | Passive wrist joint movements | Classical | Increased activation after CIMT |

Several factors may have contributed to these discrepancies such as the delay since stroke onset, the lesion topography e.g. the integrity of internal capsule (22) and, especially in stroke patients, a possible neurovascular coupling correlating the brain signal to the vascular territory of stroke (23), the impairment severity, the type of motor task performed during functional imaging, the mirror movements during functional imaging, the time-dependent changes during the rehabilitation procedure, and the type of rehabilitation (For a review see (8,24)).

Although the constraint of the unaffected arm on the one side and intensive rehabilitation on the other side are parts of most CIMT programs, the duration of the exercises varies largely between studies (5). Sub-group analyses performed in this latter Cochrane review suggested that the dosage of time practice of these two components of CIMT influences the outcome. In turn, it can be hypothesized that these two components of CIMT influence differently brain plasticity, which would partly explain the heterogeneity of functional imaging results. In a second Cochrane review, the same group confirms the strong heterogeneity among patients after CIMT (25) and the authors suggest that further trials exploring the relationship between participant characteristics and improved outcome should be performed. Proposing individual-based rehabilitation therapy requires to explore adequation between participant characteristics (e.g. lesional pattern) and specific brain activity associated with each CIMT component. This second point is exactly the purpose of our study.

## 2 Material and methods

### 2.1 Patient and controls

In this case-control study, the patient was a 32 years old right-handed woman who suffered a left capsular infarct responsible for a right hemiparesis three and a half years prior to inclusion. At the time of inclusion, the clinical examination found right sided hemiparesis, which was purely motor and consistent with an upper motoneuron lesion with brisk, diffuse osteotendinous reflexes. Her right upper limb was weak and lacked dexterity. Flexion-extension movements of wrist and fingers were possible but their active amplitude was limited to approximately 20°.

A group of healthy subjects served as a control group in determining the functional activation map associated with motor activity. We examined 10 healthy volunteers (mean age: 45.7; range: 27-62 years). All were right-handed, had a normal or corrected-to-normal vision, and had no history of neurological or psychiatric disorders. They all gave their informed consent according to the Local Ethics Committee regulation (No. AO 1280-55).

### 2.2 Experimental design

For the patient, the first therapy period involved the constraint of the non-affected arm and was followed by an intensive training after stabilization of motor performance on the Fugl-Meyer Assessment (FMA) (26) (Figure 1).

**Figure 1:**
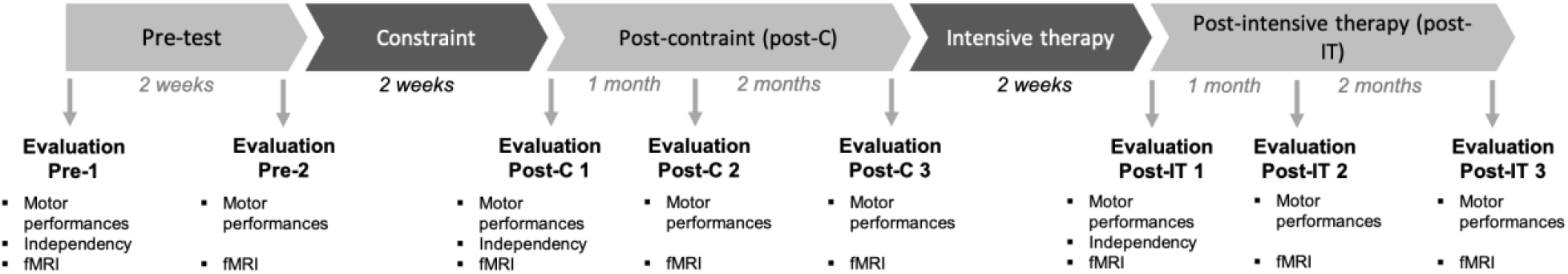
Design of the study. Experimental flowchart illustrating study design, treatments timeline and evaluations. Motor performances: Fugl-Meyer assessment scale (FMA), grip strength, box and block test, grasping task. Daily-life independency: Frenchay arm test (FAM), Functional independence measure (FIM). fMRI: functional magnetic resonance imaging.

A pre-test period was followed by the treatment program described below. During the constraint period, which lasted 2 weeks (5 days per week), the patient was instructed to wear a restrictive sling and a mitt on the non-affected upper limb during 90% of waking hours (12 hours per day). No specific exercises were prescribed during the constraint period. The second rehabilitation period was the intensive training. A six-hour daily rehabilitation program was arranged during 2 weeks (5 days per week). This consisted of physiotherapy, occupational therapy and adapted physical activities. Activities of daily living such as bathing and dressing were supervised and included in the rehabilitation time. The intensity and complexity of exercises in physiotherapy (staking objects, fine motor skills, following curves, throwing and catching a ball, precision throwing, proprioceptive neuromuscular facilitation…) and occupational therapy (building objects with pieces of wood, checkers, writing, pearl necklace, tightening or loosening nuts, darts, functional daily living exercises…) evolved with the patient’s performances according to the shaping principle. Each activity was supervised by the relevant therapist or nurse for 30 minutes followed by 30 minutes of unsupervised training. During these two rehabilitation periods, the patient was an inpatient on the neurological rehabilitation ward.

Eight time points were considered (Figure 1). At baseline, the stability of motor performance was assessed by two pre-tests (Pre) performed two weeks apart. Three post-tests were run after the constraint period (Post C): immediately, then one and two months after the constraint period. Next, three post-tests were performed after the intensive training period (Post IT): immediately, then one and two months after the intensive period.

At each time point, the motor impairment was assessed using the upper-limb section of the FMA (maximum: 66 points), a measure of grip strength, the box and block test, a three-dimensional (3D) motion analysis of a grasping task and an fMRI session were recorded. In order to assess the functional impact of each component of the CIMT, the Frenchay arm test (FAM) (27) and the functional independence measure (28) were evaluated before rehabilitation (Pre-test), after the constraint period (Post-C) and after intensive rehabilitation (Post-IT). The present study focuses on the functional imaging results; precisely, the neural changes that underlie the motor recovery induced by each CIMT component. The healthy subjects underwent the fMRI experiment in a single session.

### 2.3 The functional MRI (fMRI) experiment

We used a 1.5T MRI system (Siemens Sonata, Siemens Medical Solutions, Erlangen, Germany). A T1-weighted 3D anatomical MRI was acquired in the transverse plane (repetition time: 2120 ms, echo time: 3.9 s, field of view: 320 x 224 mm^2^, voxel size: 1×1×1 mm^3^, 384 axial slices). In order to assess the structural integrity of the internal capsule, the first MRI comprised diffusion tensor imaging sequences with the following parameters: slice number: 51, field of view: 220, slice thickness: 2.5 mm, TR: 6500 ms, TE: 86 ms, resolution: 88 x 88 (2.5 x 2.5), 24 directions, b = 1000 s/mm^2^. The fractional anisotropy asymmetry index (FA contralesional – FA ipsilesional) / (FA contralesional + FA ipsilesional) was calculated using the methodology reported by Stinear et al (29).

BOLD (Blood-oxygen-level dependent) echo planar images were collected using a T2*-weighted gradient-echo (repetition time: 2500 ms, echo time: 60 ms, field of view: 220 x 220 mm2, slice thickness: 5 mm, matrix size: 64 x 64; voxel size= 3.4 x 3.4 x 5 mm^3^). The slices were oriented parallel to the line connecting the anterior to the posterior commissure.

### 2.4 The functional MRI test task

The patient performed sequential movements opposing the thumb to the other four fingers (opposition task) with the right paretic hand. This task reflects a good level of motor recovery and has already been studied in previous fMRI study looking for neural correlates of CIMT effect (15).

The control group performed the same task with the right hand. The movement tasks were auditorily cued.

A block design was implemented with a 7-minute echo-planar imaging (EPI) run for each limb. Each run comprised seven 30-second rest periods in alternation with seven 30-second movement periods (opposition task). A foam pad was used to limit head movement during the scanning. The investigator monitored also visually the presence of mirror movements during the scanning.

### 2.5 Data analysis

#### DTI analysis

Briefly, DTI data were analyzed using Oxford Centre for Functional Magnetic resonance Imaging of the Brain (FMRIB) Software Library (FSL) (30). First, the magnetic susceptibility was corrected by Topup. The Brain Extraction Tool (BET) was used to extract the brain tissue and generate a bineary mask of the brain, created from the patient’s anatomical 3DT1 MRI. Volumes were eddy-current corrected using FMRIB Diffusion Toolbox. Finally, the tensor estimation was realized using DTIFIT to provide maps of the tensor diffusivity. Fractional anisotropy (FA) was calculated as defined by Smith et al (31). To note, FA reflects the directionality of water diffusion (from 0 (isotropic) to 1 (anosotropic)). The MedINRIA software (https://med.inria.fr) was used to extract FA in the ROI i.e. the posterior limb of the internal capsule. This ROI was delineated: i) anterior and inferior border: anterior commissure, ii) superior border: base of corona radiata, iii) posterior border: lateral ventricle, iv) medial border: thalamus, v) lateral border: putamen. The FA asymmetry index was calculated as followed: (FA_contralesional_ – FA_ipsilesional_) / (FA_contralesional_ + FA_ipsilesional_).

#### Functional MRI analysis

fMRI data were analysed using Statistical Parametric Mapping (SPM5, Wellcome Trust Centre for Neuroimaging, http://www.fil.ion.ucl.ac.uk) implemented in Matlab 7 (Mathworks, Inc, Natick, Massachusetts).

In the healthy subjects group, all the images were realigned together, spatially normalized to a standard template (MNI / ICBM) and smoothed using a Gaussian kernel with a full width at half maximum (FWHM) of 8 mm. A statistical activation map was created for each subject to contrast the images recorded during movement with those recorded during rest (movement-rest)healthy. Head motion was modelled using six regressors of no interest obtained during the realignment step. The resulting contrasts were then entered into a second-level random effects analysis (one-sample t-test). The confounding effect of age was taken into account and modelled. The resulting statistical map was corrected for Familywise Error (FWE) and thresholded at p < 0.05.

In the patient, the activations related to the movements performed with the right paretic arm were analyzed. Images were pre-processed similarly to healthy subjects’. For each time-point, a statistical activation map was computed by contrasting the images recorded during movement with those recorded during rest (movement-rest)patient. Additionally, a comparison between the two pre-tests was computed in order to verify the stability of the signal during this baseline condition. Finally, (movement-rest)patient contrasts were then entered into second-level random effects analyses (two-sample t-tests) to compare the two pre-tests, the three tests performed after the constraint period (post C), and the three tests acquired after intensive therapy (post IT) with those of the healthy subjects. The resulting statistical maps were thresholded at p < 0.001, uncorrected, and only clusters with p < 0.05 FWE corrected were reported.

#### Covariation analysis in the patient

In order to search for significant changes of brain activity across sessions, within-subject covariation analyses of brain activity and motor performance (FMA scores) were performed in patient’s data. This type of analysis is suitable to avoid a test-retest effect that cannot be ruled out in a classical factorial design. Patient’s images were realigned, spatially smoothed (FWHM of 6 mm) and spatially coregistered with the 3D-T1-weighted anatomical image. In this analysis, spatial normalization was not necessary given that all could be analysed in subject’s space.

The FMA scores of each assessment were modelled as covariates in two statistical parametric mapping (SPM) multiple regression analyses in order to search for brain regions in which a BOLD signal would have linearly changed with motor performance. The first analysis assessed the covariation between FMA scores and movement-rest contrasts issued from pre-test and the post-C sessions. A second analysis assessed the covariation between FMA scores and movement-rest contrasts issued from post-C and post-IT sessions. The resulting statistical parametric maps representing positive and negative covariations between brain activity and FMA scores were thresholded at p < 0.05 (uncorrected) at the whole brain level. Significant positive or negative correlated clusters (Small volume correction, FWE p<0.05, spatial extent threshold k>10 voxels) were selected within motor regions of interest (ROI) drawn as described below.

We used MRIcron^®^ to define a set of motor Regions Of Interst (ROIs) (Figure 2) on the basis of anatomic landmarks and standard atlases (32,33) and under the control of a neuroradiologist who was blind to the behavioural performances: (i) area M1 was defined as the posterior half of the precentral gyrus extending posteriorly to the central sulcus (17); (ii) the Premotor Cortex (PMC) encompassed the anterior half of the precentral gyrus extending anteriorly to the pre-central sulcus; (iii) the Supplementary Motor Area (SMA) was defined as the medial cortex above the cingulate gyrus, anterior to the precentral gyrus; (iv) the primary sensory cortex (S1) corresponded to the post-central gyrus; (v)(vi) the right and left part of the cerebellum were defined as the corresponding hemisphere and half of the vermis. The ROI were used for the covariation analysis Location of activations in the cerebral cortex was determined according to the stereotaxic coordinate system of the standard MNI template; location of activations in the brainstem and cerebellum was determined according to the spatially unbiased atlas template of the human cerebellum (34,35) and the three-dimensional MRI atlas of the human cerebellum in proportional stereotaxic space (36). In the expression of all results, “ipsilateral” and “contralateral” refer to the hand used to perform the task and “ipsilesional” and “contralesional” to the side of the lesion whatever the task.

**Figure 2:**
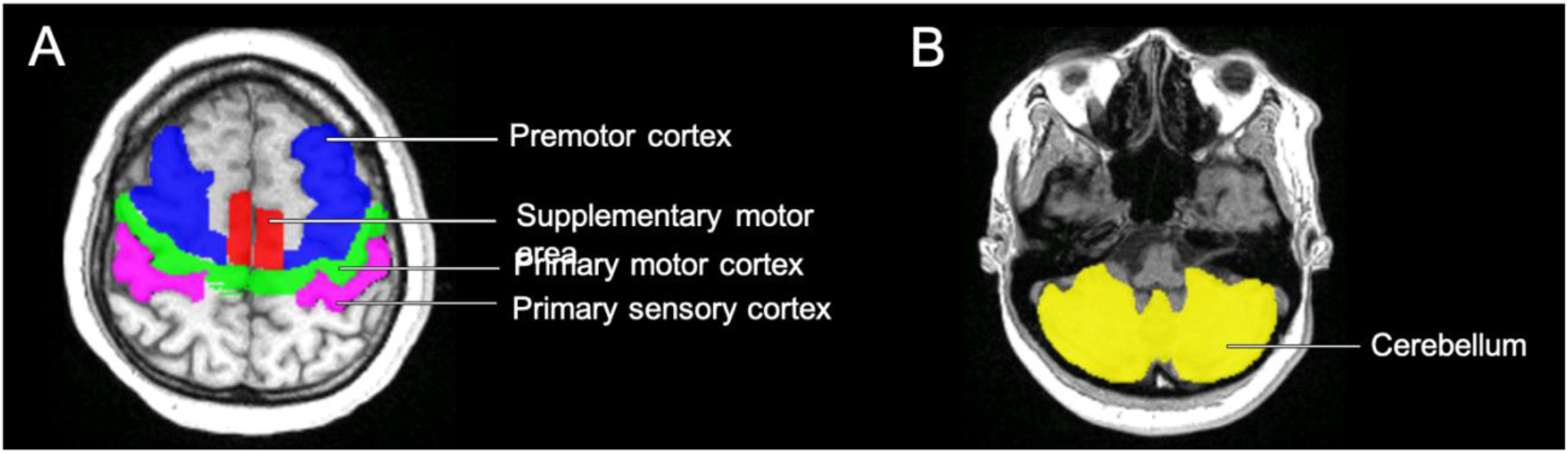
Maps of the motor Regions Of Interest (ROIs) used for the covariation analysis. A) Cortical motor regions: primary sensory cortex S1 (purple); area M1 (green); premotor cortex, PMC (blue), supplementary motor area, SMA (red). B) Cerebellum (yellow).

## 3 Results

### 3.1 Behavioral data

The motor performance as assessed by the upper-limb section of the FMA, the grip strength, and the box and block test were stable before rehabilitation (the mean score on the FMA for the two pre-tests: 28/66; mean grip strength: 9 kg; mean number of displacement on the box and block per minute:27), improved after the constraint period (mean difference between pre-tests and post-constraint: + 19 on the FMA score; + 8 kg on the grip strength; +14 displacements per minute on the box and block test), and improved again after the intensive training period (mean difference between post-constraint and post-intensive rehabilitation: + 7 on the FMA score; + 1 kg on the grip strength; +7 displacement per minute on the box and block test). The functional skills as assessed by the Frenchay arm test and the FIM also improved after constraint (+3 for the Frenchay arm test and +8 for the FIM between pre-tests and post-constraint) and after intensive rehabilitation (+1 for the Frenchay arm test and +3 for the FIM between post-constraint and post-intensive rehabilitation). The results relative to the behavioural data are shown in Table 2

**Table 2:** Progression over sessions of motor performance and independency on ADL. FMA: Fugl-Meyer assessment scale - FIM: functional independence measure

|  | Pre-tests |  |  |  | Post-constraint |  |  |  |  | Post-intensive therapy |  |  |  |
| --- | --- | --- | --- | --- | --- | --- | --- | --- | --- | --- | --- | --- | --- |
|  | Pre-1 | Pre-2 | Mean |  | Post-C1 | Post-C2 | Post-C3 | Mean |  | Post-IT1 | Post-IT2 | Post-IT3 | Mean |
| Motor performances |  |  |  |  |  |  |  |  |  |  |  |  |  |
| Upper-limb FMA, total/66 | 28 | 28 | 28 |  | 44 | 47 | 51 | 47 |  | 55 | 55 | 53 | 54 |
| Grip strength, kg | 8 | 10 | 9 |  | 16 | 17 | 17 | 17 |  | 18 | 18 | 18 | 18 |
| Box and block, per minute | 30 | 23 | 27 |  | 37 | 40 | 47 | 41 |  | 48 | 49 | 47 | 48 |
| Daily-life independency |  |  |  |  |  |  |  |  |  |  |  |  |  |
| Frenchay arm test, /5 | 1 | - | - |  | 4 | - | - | - |  | 5 | - | - | - |
| FIM, /126 | 114 | - | - |  | 122 | - | - | - |  | 125 | - | - | - |

### 3.2 Structural brain imaging

The FA asymmetry index value in the posterior limb of the internal capsule was low: 0.048. This indicates small interhemispheric differences in corticospinal tract integrity.

### 3.3 Functional brain imaging

In the healthy subjects (control group), the right-hand-opposition-task-related activity comprised the contralateral precentral gyrus, the contralateral superior temporal gyrus, the ipsilateral cerebellum and the contralateral supramarginalis gyrus (see Figure 3A and Table 3).

**Figure 3:**
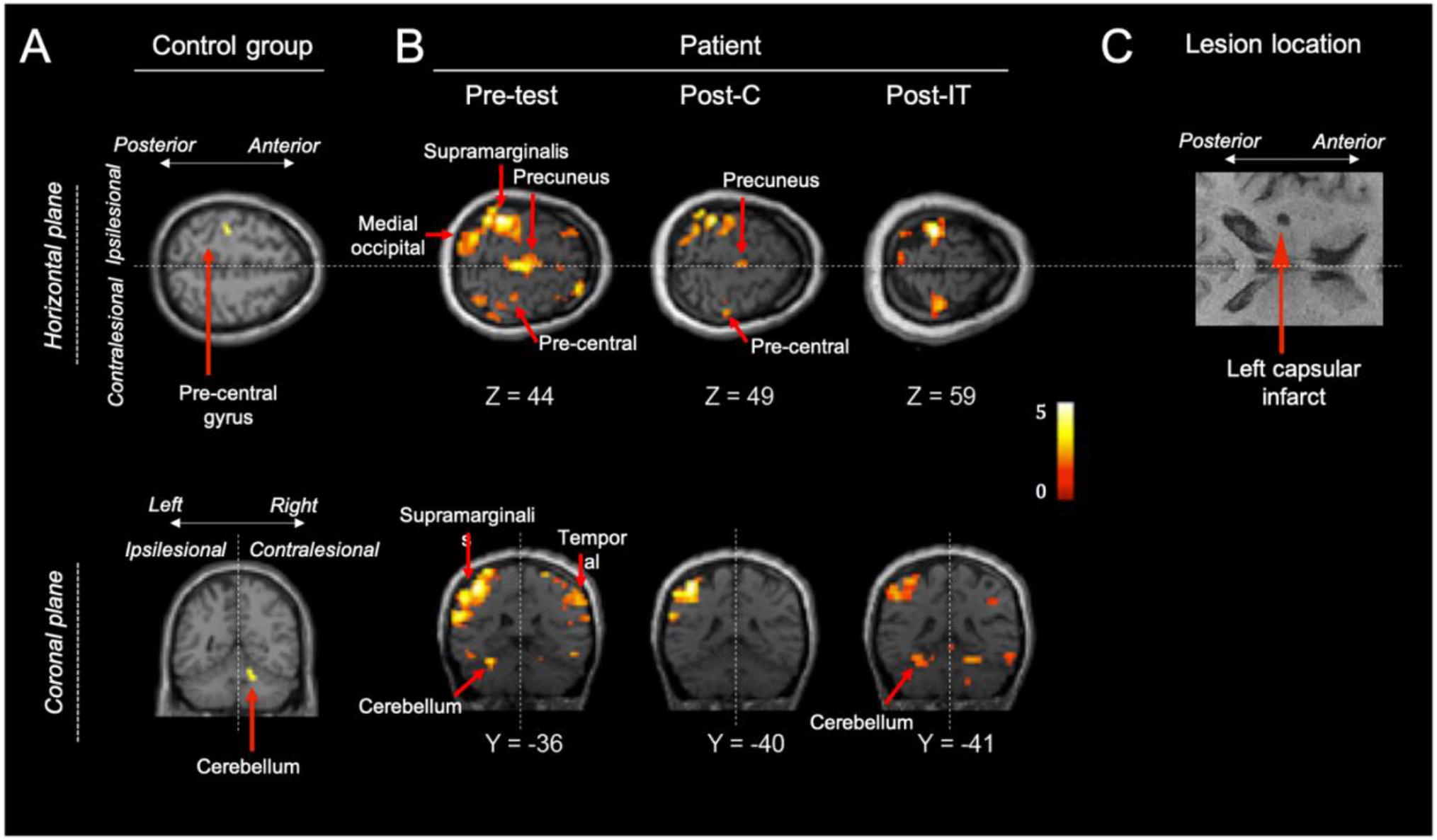
Activation pattern associated with the opposition task (illustrated on a normalized brain). A) Activation pattern associated with the finger-to-thumb opposition task performed by healthy subjects (control group) with the right hand (task vs rest). The statistical map was thresholded at p < 0.05 and corrected for Family Wise Error (FWE). B) Illustration of activation patterns associated with the finger-to-thumb opposition task (task vs rest) performed by the patient with the right paretic arm before the beginning of the therapy (2nd pre-test), after constraint (2nd test post C), and after intensive therapy (2nd test post IT). The resulting statistical map was thresholded at p < 0.001; functional images are not spatially normalized and are displayed on the patient’s structural MRI. C) Structural T1-weighted image showing the lesion location in the posterior limb of the left internal capsule. The left hemisphere is displayed on the left side.

**Table 3:**
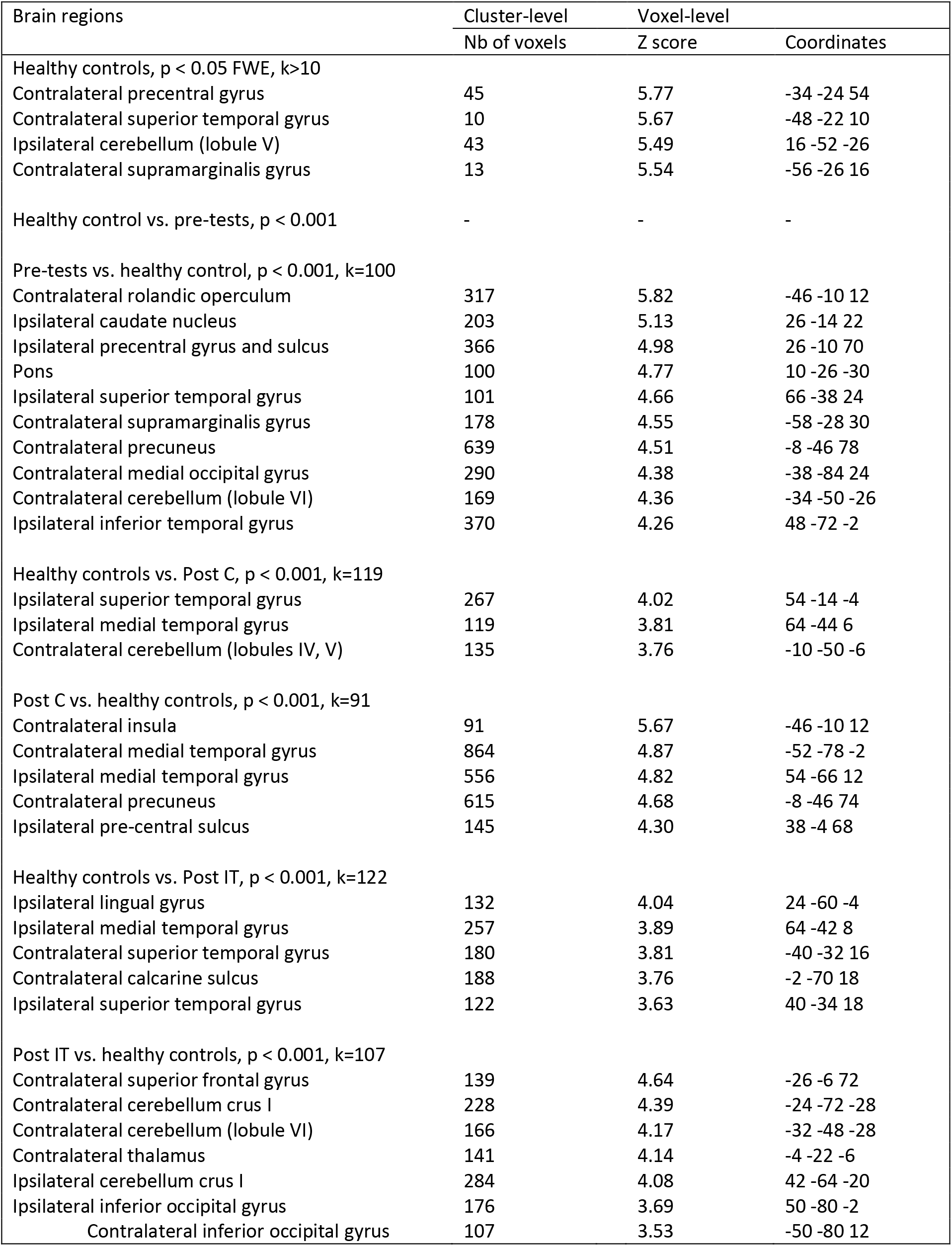
Functional activation associated with the finger opposition task. FWE: Familywise Error - post C: post constraint - post IT: post intensive training - -: No activation found - “Ipsilateral” and “contralateral” refer to the hand used to perform the task-Comparisons between Healthy subjects and Patient are reported at p<0.001 uncorrected at the voxel level, extent cluster value *k* is set in order to select clusters reaching significance of p <0.05 FWE corrected.

The patient performed the opposition task in a reproducible manner without visible mirror movement of the other limbs.

The stability of the two pre-tests periods was assessed and only two regions were found significantly more activated during the 2nd pre-rehab period than in the 1st pre-rehab period. These regions were identified as the contralateral (ipsilesional) post-central gyrus (−40 −24 66, p<0.05 corr, KE > 10) and the ipsilateral (contralesional) inferior frontal gyrus (50 38 −6, p<0.05 corr, KE>10). No regions were significantly activated during the 1^st^ period when compared to the 2^nd^ period.

When compared to the control group, the opposition task performed by the patient with the paretic hand during the pre-tests (Figure 3B, “Pre-test”) recruited a larger network including the pons and brain areas both in the contralateral (ipsilesional) hemisphere (rolandic operculum, supramarginalis gyrus, precuneus, medial occipital gyrus and lobule VI of the cerebellum) and in the ipsilateral (contralesional) hemisphere (caudate nucleus, precentral gyrus and sulcus, superior temporal gyrus, inferior temporal gyrus). After the constraint period, substantial differences in the pattern of activation were also observed as compared to healthy controls (Figure 3B, “Post-C”): the contralateral (ipsilesional) insula, the bilateral medial temporal gyrus, the contralateral precuneus, the ipsilateral (contralesional) pre-central sulcus were more active in the patient than in the controls. The ipsilateral superior temporal gyrus, the ipsilateral medial temporal gyrus, the contralateral cerebellum (lobule IV and V) were more active in the opposite contrast (healthy controls vs. post C). Constraint-therapy-related activation was inversely correlated with FMA improvement in bilateral M1 and S1, ipsilateral (contralesional) PMC, bilateral SMA, and the contralateral (ipsilesional) cerebellum (see table 4). No region was found to be positively correlated with FMA. With intensive-therapy-related activation, a positive correlation with FMA was found in bilateral PMC and the cerebellum, bilaterally, whereas an inverse correlation was found only in the contralateral S1.

**Table 4:** Brain activity correlated with Fugl-Meyer Assessment (FMA) score across sessions. FMA: Fugl-Meyer Assessment; M1: primary motor cortex; S1: primary somato-sensory cortex; SMA: supplementary motor area; PM: pre-motor cortex; IT: intensive training.; ipsilateral and contralateral refers to the hand used to perform the task. Significant clusters after Small Volume Correction (FWE p<0.05, ke =10). Identification is based on motor ROI (cf Methods) drawn in subject’s space (i.e. no coordinates available).

| Regions of interest | Voxels | Z score |
| --- | --- | --- |
| Positive correlation between constraint-therapy-related activation and FMA | No significant voxel |  |
| Negative correlation between constraint-therapy-related activation and FMA |  |  |
| Ipsilateral M1 | 33 | 3.27 |
| Contralateral M1 | 12 | 2.84 |
| Ipsilateral S1 | 30 | 3.27 |
| Contralateral S1 (2 clusters) |  |  |
| Cluster 1 | 10 | 3.11 |
| Cluster 2 | 17 | 2.57 |
| Ipsilateral PM (2 clusters) |  |  |
| Cluster 1 | 71 | 3.18 |
| Cluster 2 | 13 | 2.44 |
| Ipsilateral SMA | 13 | 2.65 |
| Contralateral SMA (2 clusters) |  |  |
| Cluster 1 | 10 | 2.93 |
| Cluster 2 | 13 | 2.83 |
| Contralateral cerebellum (lobules III and IV) | 110 | 3.37 |
| Positive correlation between IT-therapy-related activation and FMA |  |  |
| Ipsilateral PM | 73 | 3.18 |
| Contralateral PM | 31 | 3.19 |
| Ipsilateral cerebellum (lobules V and VI) | 210 | 3.54 |
| Contralateral cerebellum (2 clusters) |  |  |
| Cluster 1 (lobules Crus I and II) | 183 | 3.53 |
| Cluster 2 (lobule VIII B) | 14 | 2.55 |
| Negative correlation between IT-therapy-related activation and FMA |  |  |
| Contralateral S1 | 23 | 2.73 |

After the intensive therapy period (Figure 3B, “Post-IT”), the superior frontal gyrus, the thalamus and the cerebellum (lobule crus I and lobule VI) within the contralateral (ipsilesional) hemisphere and the inferior occipital gyrus bilaterally were found more active in the patient as compared to the healthy controls. The opposite contrast (healthy controls vs. post C) revealed an activation of the lingual, medial temporal gyrus and superior temporal gyrus within the ipsilateral (contralesional) hemisphere, the superior temporal gyrus and the calcarine sulcus within the contralateral (ispilesional) hemisphere.

## 4 Discussion

In this present study, we investigate the brain reorganization after the two components of CIMT - i.e. after the constraint phase and after the intensive therapy. Functional motor performances were improved by each component of the CIMT, assessed by gold standard clinical scales (Frenchay arm test or *FMA* and the functional independence measure or FIM). Constraint-therapy-related improvement was correlated with a decrease of cerebral activity in sensory-motor regions (M1, S1, SMA) of both the affected and the non-affected hemispheres. Intensive-therapy-related improvement was correlated with the recruitment of pre-motor cortices and cerebellum in both hemispheres. Finally, three months after the intensive therapy, the patient still exhibited an increased activity ipsilesional thalamus and cerebellum.

Constraint-therapy-related improvement was correlated with a decrease of cerebral activity in sensory-motor regions ( of both the affected and the non-affected hemispheres. Intensive-therapy-related improvement was correlated with the recruitment of pre-motor cortices and cerebellum in both hemispheres.

The present longitudinal functional imaging study was developed to investigate step-by-step the brain plasticity mechanisms elicited by two components of CIMT −constraint and intensive training– in a post-stroke patient. Although at a chronical stage (the study began more than 3 years after stroke), the patient showed a remarkable improvement after each component of CIMT. The gain on the upper-limb section of the FMA, used as a covariate to study neural reorganization associated with CIMT effect, is clinically significant (+19 between pre-tests and post-constraint and + 7 between post-constraint and post-intensive rehabilitation). According to Sanford et al (37), a difference on the upper-limb section of the FMA is reliable when the difference is above 3.6. The first step of this study was designed to explore the normal pattern of brain areas activated during a finger opposition task. Our results confirmed the classical implication of the contralateral primary motor cortex and the ipsilateral cerebellum during a finger motor task (38). Three other areas were also significantly more activated during the task than during rest: the contralateral superior temporal gyrus, the contralateral supramarginalis gyrus. These regions are less consistently observed; their activation reflects the specificity and complexity, beyond motor execution, of the opposition motor task employed here.

The left superior temporal gyrus is involved in auditory integration and language processing. One obvious explanation for its activation in the present study is the auditory cue present during the movement periods and absent during the control rest periods.

The opposition of the thumb to the four other fingers comprised a spatio-temporal sequence that might explain the activation of the supramarginalis gyrus (SMG) known to be implicated in attention (39) and in temporal synchronization (40). Another hypothesis for this activation is the mental imagery process that accompanies finger movements. Indeed, the inferior parietal area has been shown to be activated during mental imagery tasks (41).

Finally, no activation was detected in the basal ganglia in the healthy subjects. This may be linked to a lack of sensitivity of the imaging technique (42) and/or to the threshold chosen for the study. It should be noted that a bilateral activation of the basal ganglia can be objectified in healthy subjects when an uncorrected threshold is applied in our analysis (not shown).

The comparison of finger opposition-related activities between the patient and the healthy controls led us to study brain plasticity after a circumscribed lesion of the descending motor pathway. The long delay since stroke onset is compatible with many neuroplasticity mechanisms such as motor map enlargement, changing the activity of the neurons (e.g. strengthening) within a specific area, network-related function changes (for a review, see (10,43)). As stated in the introduction, our functional imaging study enables us to discuss mainly the vicariation versus the interhemispheric competition models. In addition to the network of brain areas that subserve the finger opposition task in our healthy control group, many motor and non-motor regions of both hemispheres as well as sub-cortical regions were recruited or overactivated in the patient during the pre-tests. As previously reported in recovered hemiparetic striato-capsular stroke patients (44–46), we observed a larger and more bilateral pattern of activation during finger movement performed by our patient as compared to the healthy controls. The spatial recruitment of additional sensori-motor cortices is likely to reflect an effortful action (47). The recruitment of non-motor regions could also reflect an increased attention when the finger opposition task is performed with the paretic arm as compared with the non-paretic one.

The FA asymmetry index was very low (0.048), far lower than the 0.25 cut-off value calculated in the prediction recovery potential (PREP) algorithm for chronic stroke patients (29). This indicates a relative preservation of the affected internal capsule and is consistent with an efficient ipsilesional descending pathway that conducts the motor output signal.

As mentioned earlier in the discussion, in this intervention study, an order effect and a test-retest effect cannot be ruled out by a classical factorial design. To circumvent this potential bias, a covariation analysis was performed to search for specific brain areas associated with motor recovery after constraint and after intensive training. The results showed clearly two different changes in the patterns of activation during a motor task that resulted from constraint versus intensive rehabilitation.

We showed that the activity in the sensory-motor regions of both the affected and the unaffected hemispheres was inversely correlated with the arm function recovery induced by constraint-therapy. This important reduction of activity is not intuitive and some authors have reported opposite results. For instance, Laible et al. (48) showed an inverse correlation between motor recovery after CIMT and activation within the ipsilesional S1. Similarly, Könönen et al. (14) showed that increased activation within the sensorimotor areas was greater amongst patients whose motor behavior improved after CIMT. Alternatively, results observed in the present study may be interpreted as a decrease of attention resources in performing the same task after the constraint period, which is similar to what is observed in spontaneous motor recovery (49,50). This activity reduction was not homogeneous. We observed an inverse correlation between the BOLD signal and motor recovery in the ipsilateral (contralesional) PMC but not in the contralateral (ipsilesional) PMC. The premotor cortex of the unaffected hemisphere has been shown to play a critical role during motor recovery (13,51). In this latter study, Johansen-Berg et al. showed that the inhibition of the ipsilateral premotor cortex by transcranial magnetic stimulation (TMS) slowed the reaction-time during finger movements in patients with right hemiparesis but not in healthy subjects. It remains unclear whether the contralesional premotor cortex participates to spontaneous recovery or reflects a maladaptative plasticity through an imbalance of interhemispheric inhibition (cf. Introduction). The latter hypothesis postulating that the ipsilateral cortex influences adversely spontaneous recovery after stroke through an interhemispheric mutual modulation of homologous areas has already been demonstrated in spatial neglect (52) and upper-limb hemiparesis (53). In the present study, the decrease of activity in the ipsilateral (contralesional) PMC as a function of motor recovery could reflect the restoration of the inter-hemispheric balance for motor control induced by constraint.

We have previously reported the effects of constraint and intensive rehabilitation on kinematic analysis and showed that grasping was selectively improved after intensive training (54). We postulated that intensive therapy could improve upper-limb prehension recovery via an enlargement of the ipsilesional M1 cortical hand representation, which is in accordance with other motor mapping experiences in monkeys (2) and human (14,55). Moreover, the enlargement of M1 activity is also known to be implicated in complex motor skill learning in healthy subjects (56). In this latter fMRI study, the training of a complex sequence of finger-to-thumb opposition movements involved the enlargement of contralateral M1. However, we did not find a significant correlation between the activation within the primary motor cortex of the affected hemisphere and the motor improvement after intensive training. At least three explanations for this negative result may be put forward: (i) Activation in the primary motor cortex (M1) of the affected hemisphere had already reached normal functioning before intensive training. This hypothesis is supported by the absence of significant difference in this region between post intensive training and healthy controls (cf Table 3). (ii) The enlargement of recruited neurons in the motor cortex might not be easily accessible by functional imaging and could be better assessed by TMS. (iii) The implication of the primary motor cortex in previous CIMT functional imaging studies differs from one study to another and may interact with stroke chronicity size and lesion topography (9,17,29). Another potential mechanism, not assessed in the current study, is a modification of overlapping representations of adjacent joints induced by CIMT. This has been demonstrated in a previous fMRI study in which brain activity of wrist, elbow and ankle movements were assessed before and after CIMT (19).

Here, the motor improvement after intensive training was correlated with an increased activity in the cerebellum and the PMC bilaterally. Similarly, in a previous fMRI study performed on seven chronic hemiparetic patients, Johansen-Berg et al. (13) have shown that motor improvement after a two-week rehabilitation period was correlated with an increased activity in the cerebellum bilaterally and in the motor and pre-motor cortices of the affected hemisphere. In both studies, the upper-limb motor improvement induced by rehabilitation after stroke was significantly correlated with an increased activity within the cerebellum and the PMC. It should be noted that increased activity in the PMC after CIMT has been also reported by Zhao et al. (21) in another fMRI study. In this latter study, activation changes were obtained in a pre-post comparison. Interestingly, both PMC and cerebellum are known to be involved in motor skill learning (57), especially at the early stages (58), and in bi-manual skill acquisition (34,59). The cerebellum is also a recognised region involved in sensori-motor adaptation, as demonstrated by functional imaging (60) and lesioned studies showing that patients with cerebellar lesions do not adapt to prisms as well as controls, although they still can learn (61,62). Hence, it can be hypothesized that, in the present study, the improvement of the upper-limb function after intensive training was mediated by the recruitment of remote motor regions involved in motor skill learning, bimanual skill acquisition, and sensori-motor adaptation. Interestingly, this hypothesis could be tested if the fMRI paradigm used in the present study is repeated multiple times in healthy subjects.

Three main limitations must be pointed out: (i) in this case-control study, the generalization is necessarily limited to stroke patients having a similar profile; (ii) an order effect cannot be ruled out because the constraint period took place before intensive rehabilitation; (iii) despite our efforts to control as much as possible the amplitude, the hand position, and the movement speed, brain activity changes could reflect modifications in the way the task was performed by the patient throughout the whole experiment as compared with healthy controls. To avoid the latter potential bias, we controlled the position of the hand as well as the amplitude and the speed of the finger movement. These visual checks during MRI acquisitions did not report mirror movements of the unaffected limb though infra-visible movements cannot be excluded in the absence of an electromyogram recording. Difference in the speed and force of the movement between the patient and the controls cannot be excluded and could have participated to differences observed in the analysis.

## 5 Conclusion

Altogether, these results illustrate the complexity of investigating the way by which a given intervention is able to modulate brain plasticity in the domain of neurological rehabilitation. Our results show two different functional brain reorganizations underlying the effects of intensive rehabilitation and constraint and argue for a “two brain plasticity” mechanisms induced by CIMT. These results may partly explain the important heterogeneous results of previous CIMT functional imaging studies and be used to fit the rehabilitation of prehension to a more theory-driven approach.

## 7 Author Contributions

SB: acquisition of data, analysis and interpretation, critical revision of the manuscript for important intellectual content. DI, JR, JCC: acquisition of data and analysis, critical revision of the manuscript for important intellectual content. BM, LA, AM, PR, LD: interpretation and critical of the manuscript for important intellectual content. FC: data analysis and critical revision of the manuscript for important intellectual content. GR, YR, DB, JL: study concept and design interpretation, critical revision of the manuscript for important intellectual content. MBG: critical revision of the manuscript for important intellectual content. All authors read and approved the final manuscript.

## 9 Acknowledgments

We thank Jean Iwaz (Hospices Civils de Lyon) for the thorough revision of the final drafts of the manuscript.

